# NS-Forest: A machine learning method for the objective identification of minimum marker gene combinations for cell type determination from single cell RNA sequencing

**DOI:** 10.1101/2020.09.23.308932

**Authors:** Brian Aevermann, Yun Zhang, Mark Novotny, Trygve Bakken, Jeremy Miller, Rebecca Hodge, Boudewijn Lelieveldt, Ed Lein, Richard H. Scheuermann

## Abstract

Single cell genomics is rapidly advancing our knowledge of cell phenotypic types and states. Driven by single cell/nucleus RNA sequencing (scRNA-seq) data, comprehensive atlas projects covering a wide range of organisms and tissues are currently underway. As a result, it is critical that the cell transcriptional phenotypes discovered are defined and disseminated in a consistent and concise manner. Molecular biomarkers have historically played an important role in biological research, from defining immune cell-types by surface protein expression to defining diseases by molecular drivers. Here we describe a machine learning-based marker gene selection algorithm, NS-Forest version 2.0, which leverages the non-linear attributes of random forest feature selection and a binary expression scoring approach to discover the minimal marker gene expression combinations that precisely captures the cell type identity represented in the complete scRNA-seq transcriptional profiles. The marker genes selected provide a barcode of the necessary and sufficient characteristics for semantic cell type definition and serve as useful tools for downstream biological investigation. The use of NS-Forest to identify marker genes for human brain middle temporal gyrus cell types reveals the importance of cell signaling and non-coding RNAs in neuronal cell type identity.

## Introduction

Cells are the fundamental functional units of life. In multicellular organisms, different cell types play different physiological roles in the body. The identity and function of a cell - the cell phenotype - is dictated by the subset of genes/proteins expressed in that cell at any given point in time. Abnormalities in this expressed genome are disorders that form the physical basis of disease (1) Thus, understanding normal and abnormal cellular phenotypes is key for diagnosing disease and for identifying therapeutic targets.

Single cell transcriptomic technologies that measure cell phenotypes using single cell/single nucleus RNA sequencing (scRNA-seq) are revolutionizing cellular biology. The expression profiles produced by these technologies can be used to define cell types and their states. Numerous atlas projects designed to provide a comprehensive enumeration of normal cell types and states are currently underway, including the Human Cell Atlas (2), California Institute for Regenerative Medicine (CIRM) (3-5), LungMAP (6), Pancreas atlas (7), Heart atlas (8), and NIH Brain initiative (9). By leveraging these atlases of normal cell types defined from healthy patients as references, the role of expression deviations in disease are being investigated (10-12).

These projects rely primarily upon two scRNA-seq single cell technologies: Droplet based technologies (13) or FACS sorting followed by Smart-Seq library preparation (14). In the standard data processing pipeline, the raw sequencing reads are processed using reference-based alignment, transcript reconstruction, and expression level estimation. From the expression matrix, the typical downstream analysis workflows produce a set of gene expression data clusters representing cells/nuclei with similar expression patterns. We interpret these distinct transcriptional profiles to represent distinct cell phenotypes, which include canonical cell types and distinct cell states, that have achieved a state of equilibrium. This is in contrast to transitional trajectories in developing tissues and populations in the process of responding to perturbations that can show gradual expression pattern changes between discrete cell phenotypes (15-17).

Despite the incredible promise of single cell transcriptomic analysis for identifying and quantifying known and novel cell types, the cell type clusters and their transcriptional phenotypes are not being formalized in a standardized way to ensure dissemination is in accordance with FAIR principles (18). One approach for formalizing knowledge representation and dissemination is to use the sematic framework provided by biomedical ontologies. For cell phenotypes defined by single cell transcriptomics, the Cell Ontology (CL) is an established biomedical ontology already applying ontological methodologies that could be used to address FAIR-compliant cell phenotype dissemination (19-22). With the rapid expansion in both datasets and cell phenotypes being defined using scRNA-seq, the challenge will be to make the generation of these semantic knowledge representations scalable.

To develop this scalable dissemination solution, we propose to define cell type phenotypes based on the minimum combination of necessary and sufficient features that capture cell type identity and uniquely characterize a discrete cell phenotype. In the case of cell types identified by scRNA-seq experiments, these features correspond to the combination of differentially expressed marker genes unique to a given gene expression cluster. Historically, marker gene expression, especially at the protein level, has been an essential tool to connect cell type identity with defining cell type functional characteristics. For example, the classical markers CD19 and CD3 have been used extensively to differentiate between B cells and T cells, while within the T cell population CD3 and CD4 and CD8 are used to further separate helper and cytotoxic types (23). In neurology, SLC17A7 and GAD1 are well-known markers for excitatory (glutamatergic) and inhibitory (GABAergic) neuron types, respectively (24).

In the case of scRNA-seq expression clusters, an optimal cell type marker would be a cellular feature or unique combination of features that provides high sensitivity and high specificity for cell type classification. The ideal scenario would be to have a marker gene that is expressed at high levels in all individual cells of a given cell type and not expressed at all in any cell of any other cell type. We refer to this phenomenon as a binary expression pattern. Finding markers with this binary expression pattern can be quite useful for downstream experimental validation and investigation using technologies such as multiplex fluorescence in situ hybridization (mFISH), quantitative PCR, or flow cytometry. However, in complex tissues this ideal scenario is rarely observed. Candidate marker genes are often expressed at high levels in the target cluster and lower levels in off-target clusters. We refer to these markers as quantitative markers as their discriminatory power is derived from specific expression level cutoffs, which are dependent on the sensitivity of the assay being performed. Or a single binary marker may be expressed in multiple related cell types.

Though similar in concept, determining markers from cell type clusters is different from differential expression analysis (DE) in that DE analysis evaluates each gene for expression level variation between groups, whereas marker genes are tested for their classification power. The most common scRNA-seq analysis tools - Seurat (16) and Scanpy (17) – handle gene selection in a similar fashion. After cluster analysis, genes are evaluated by comparing expression in cells in a target cluster versus expression in all other cells using, for example, the Wilcoxon Rank Sum test, which produces a gene list that can be ranked by adjusted p-value. However, while the best marker genes for discriminating and defining a cell type cluster are often found among the differentially expressed genes, their utility in defining a cell type is not apparent from either their p-value rank nor their fold difference in expression. Furthermore, DE analysis tests genes individually, while defining cell types often requires the combined contribution of sets of marker genes.

Here we describe Necessary and Sufficient Forest (NS-Forest) version 2.0, which leverages the non-linear attributes of random forest feature selection and Binary Expression Score ranking to discover marker gene combinations that can be used to both define cell type phenotypes and in downstream biological investigations. The initial version of NS-forest was based on a simple approach in which feature selection by Random Forest machine learning was used to discover potential marker genes (21). Here we describe user community-driven improvements upon this original methodology. NS-Forest version 2.0 is available at https://github.com/JCVenterInstitute/NSForest under an open source MIT license.

## Results

### User driven development of NS-Forest

NS-Forest version 2.0 was developed in close collaboration with the brain cell user community. The primary goal was to optimize the NS-Forest method in order to discover marker genes that can both uniquely define the cell type phenotype and aid in their downstream experimental investigation. In order to accomplish this, several major changes were made to NS-Forest version 1 (**Table 1**). First, negative markers were removed by implementing a positive expression level filter. A negative marker is not expressed in the target cluster while having expression in off-target clusters. These markers are not optimal for many downstream assays or definitional purposes. These genes are now filtered out by applying a cluster median expression threshold. The default setting is zero; however, this can be changed to enrich for genes at varying expression levels (**Figure 1C**).

**Table 1:** Major changes between NS-Forest version 1.3 and version 2.0

| Workflow step | NS-Forest v1.3 | NS-Forest v2.0 |
| --- | --- | --- |
| Feature Selection (Figure1A/B) | Random Forest (RF) driven feature selection | No changes |
| Feature Ranking (Figure1C) | Used RF ranking | Filtering of negative markers |
| Feature Ranking (Figure1D/E) | NA | Binary score reranking |
| Minimum feature determination (Figure1F) | Gene expression criteria determined by median cluster expression | Gene expression criteria determined by Decision Tree split |
| Minimum feature determination (Figure1G) | Calculate F-score | Calculate Fbeta-score |

**Figure 1:**
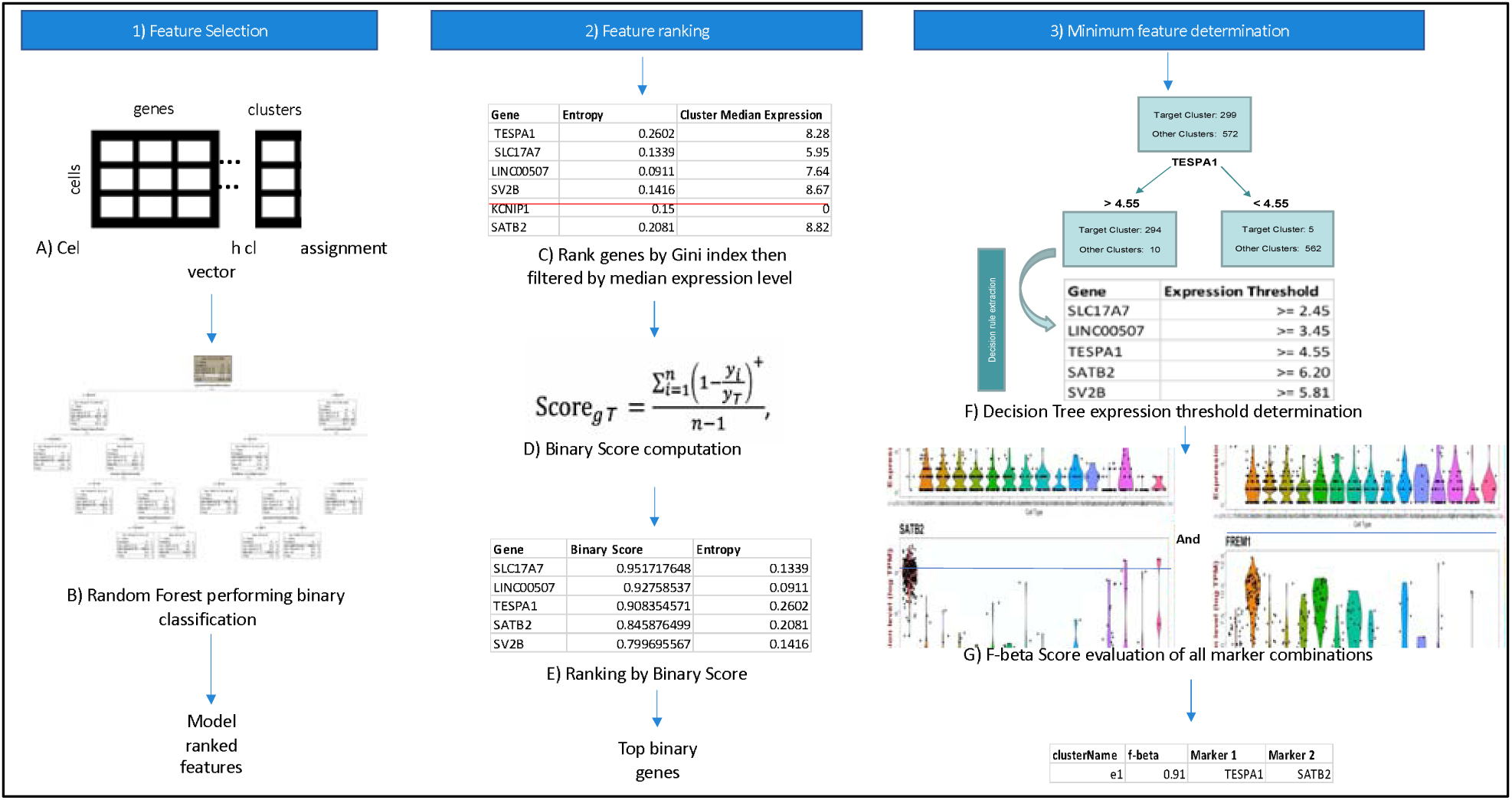
NS-Forest version 2.0 workflow. The method begins with a cell by gene matrix and cluster assignments. These are used to generate binary classification models using Random Forest. Features are extracted from the model and ranked by Gini index. Top features are filtered by expression level before being ranked by binary expression. Decision point expression level cutoffs are then determined for the most binary features and F-beta score used as an objective function to evaluate the discriminatory power of all combinations.

Next, the way genes are ranked after Random Forest (RF) selection was refined. Genes selected by RF have an expression level split point between target and off-target clusters. Often the genes selected discriminate based on a specific value of expression resulting in quantitative expression markers. As will be demonstrated below, these quantitative markers are good for classification but are less useful in many downstream biological assays. To address this issue, we optimized version 2.0 for the selection of binary expression markers. Binary expression markers are characterized by having expression within the target cell type while being expressed at low or negligible levels in other cell types. We accomplished this by developing a new Binary Expression Score metric with subsequent re-ranking based on this score after random forest feature determination (**Figure 1D/E**).

Lastly, the marker gene evaluation framework was redesigned. In early NS-Forest versions, the top ranked genes were evaluated by unweighted F-1 score in an additive fashion. First, each of the top genes were tested individually and the best gene then removed from the list. Next all pairings with the previously determined gene were tested to find an improvement over the previous individual gene F-score. This additive process continued until the F-score plateaued or the selected number of top rank genes were all tested. In the new version of NS-Forest, all combinations of the selected top ranked genes are tested by weighted F-beta score. The F-beta score contains a weighting term, beta, that allows for emphasizing either precision or recall. By weighting toward precision, zero inflation (drop-out) can be controlled, which is a known technical artifact with scRNA-seq data. These adjustments result in better final marker gene combinations given the known limitations of scRNA-seq analysis (**Figure 1F/G)**.

### Performance Testing of Binary Scoring Approach

Simulation testing of the NS-Forest Binary Expression Score was performed to evaluate re-ranking behavior. First, anticipated marker gene expression patterns were themselves ranked by order of hypothetical preference (**Figure 2A)**. The highest preference was given to a marker gene that shows a binary expression pattern and is only expressed in the target cluster (**Figure 2A(a)/2B**). Next, preference is given to a marker gene that shows binary expression and is only expressed in the target cluster and a limited number of off-target clusters (**Figure 2A(b))**. This is followed by quantitative markers which have a high expression in the target cluster and lower expression in off-target clusters (**Figure 2A(c)/2C**) or high expression in the target cluster and a limited number of off-target clusters (**Figure 2A(d)**). The least preferred pattern is when the marker is expressed at only slightly different levels between the target and off-target clusters (**Figure 2A(e)/2D**).

**Figure 2:**
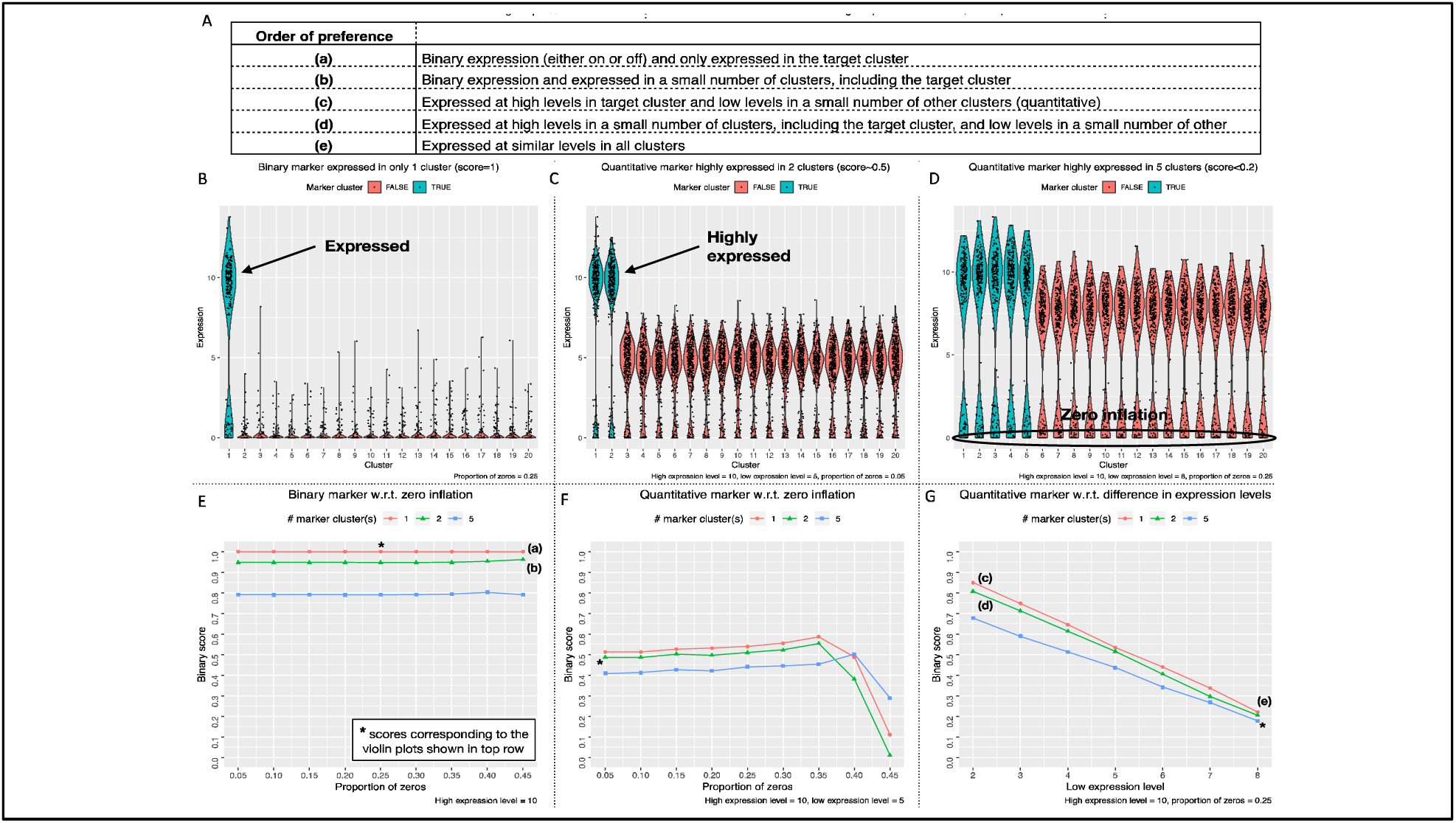
Performance testing of Binary Expression Score. In panel A) Possible marker gene expression patterns were ranked by order of preference. Violin plots showing three different expression scenarios: panel B) binary expression only in the target cluster, panel C) quantitative expression with high expression in the target cluster and one other cluster and large differences in expression in the other off-target clusters, panel D) quantitative expression with high expression in the target cluster and four other cluster, small differences in expression in the other off-target clusters, and higher levels of zero inflation. Below, line graphs show the full range of tested simulations from which the above example violin plots are taken. For panels E-G there were three defined test cases: the red where there is one cluster with high expression of the marker gene, green where there are two clusters with high expression of the marker gene, and lastly blue where 5 clusters have high expression of the marker gene. Panel E) gives simulation testing of the Binary Expression Score increasing the proportion of zeros while maintaining off target expression at zero. Panel F) off-target clusters were given moderate levels of expression while the proportion of zeros was increased. In panel G) expression levels were varied in all off-target clusters from low (2) to high expression (8).

Simulations varying the binary expression pattern and level of zero inflation (**Figure 2E**) were then performed. First, the ideal scenario of binary expression, as described above, produced a simulated Binary Expression Score of 1.0 (**Figure 2E** red). When the candidate marker gene was expressed in one (**Figure 2E** green) or four (**Figure 2E** blue) off-target clusters, the Binary Expression Score decreased to 0.95 and 0.90, respectively. In addition, these scores were robust to high zero inflation proportions, demonstrating no decrease in Binary Expression Score up to 45% zero values.

Next, quantitative marker expression patterns was added to the simulation (**Figures 2F & G**) by varying the number of off-target clusters with high expression levels and adding moderate expression to the remaining off-target clusters. In all cases in which quantitative difference in expression are simulated, the Binary Expression Scores are substantially reduced (**Figures 2F)**. In the best case, where only the target cluster has high expression and the off-target clusters have moderate expression, the Binary Expression Score was 0.52. Further Binary Expression Score reductions are found when the high expression levels are present in additional off-target clusters. Adjusting the level of zero inflation for these scenarios showed that these Binary Expression Scores were also robust to increasing zero inflation levels until they drop dramatically after 35% zero values.

Finally, simulations were performed to again test how a high-expressing marker is effected by the addition of 1 or 4 high expressing off-target clusters together with increasing expression levels in the remaining off-target clusters from low (2) to high (8) expression (**Figures 2G**). With the remaining off-target clusters held at low expression levels, these three scenarios returned high Binary Expression Scores [0.7-0.85], but these Binary Expression Scores quickly decreased with increasing levels of off-target expression. For example, when the off-target expression level was set to 6, all three high-expressing off-target scenarios returned Binary Expression Scores below 0.5. In the worst case, where the candidate marker has relatively high expression in all off-target clusters, the Binary Expression Score was less than 0.2.

These simulations demonstrate that the Binary Expression Score value produced by the algorithm recapitulates the preferred expression pattern ranking order (**Figure 2A)**. In all simulations tested, the Binary Expression Score decreases with the addition of marker expression in off-target clusters and were robust to zero inflation.

### Marker Gene Comparison Between NS-Forest Versions

To evaluate the differences in results between NS-Forest v1.3 and v2.0, we analyzed marker genes selected for cell type clusters generated from single nuclei transcriptomes prepared from all layers (1-6) of the human middle temporal gyrus (MTG) obtained from postmortem and surgically resected samples (**sTables 1-3**). For this dataset, three broad classes of cells were identified: excitatory neurons (10,708 cells), inhibitory neurons (4,297 cells), and non-neuronal cells (923 cells). These nuclei were clustered iteratively by first clustering into the larger groups, followed by subsequent re-clustering within each group until 75 putative cell types were found (25). From left to right of the hierarchical clustering of clusters shown at the top of both heatmaps there are 46 inhibitory, 23 excitatory, and 6 non-neuronal types (**Figure 3)**. Subsequent figures investigating these cell type clusters are ordered by these taxonomic relationship. In Figure 3, the 155 marker genes determined by NS-Forest version 1.3 and the 157 marker genes determined by version 2.0 are the rows while the clusters are columns where the row normalized expression level is reflected in the color gradient of high in red to low in blue/white. The diagonal corresponds to the marker gene set for each cluster type. From the heatmaps it is clear that the binary expression has dramatically improved between NS-Forest version 1.3 and version 2.0 as the diagonal contains more genes with red or bright yellow levels of expression and off-diagonal expression levels are more blue (closer to 0).

**Figure 3:**
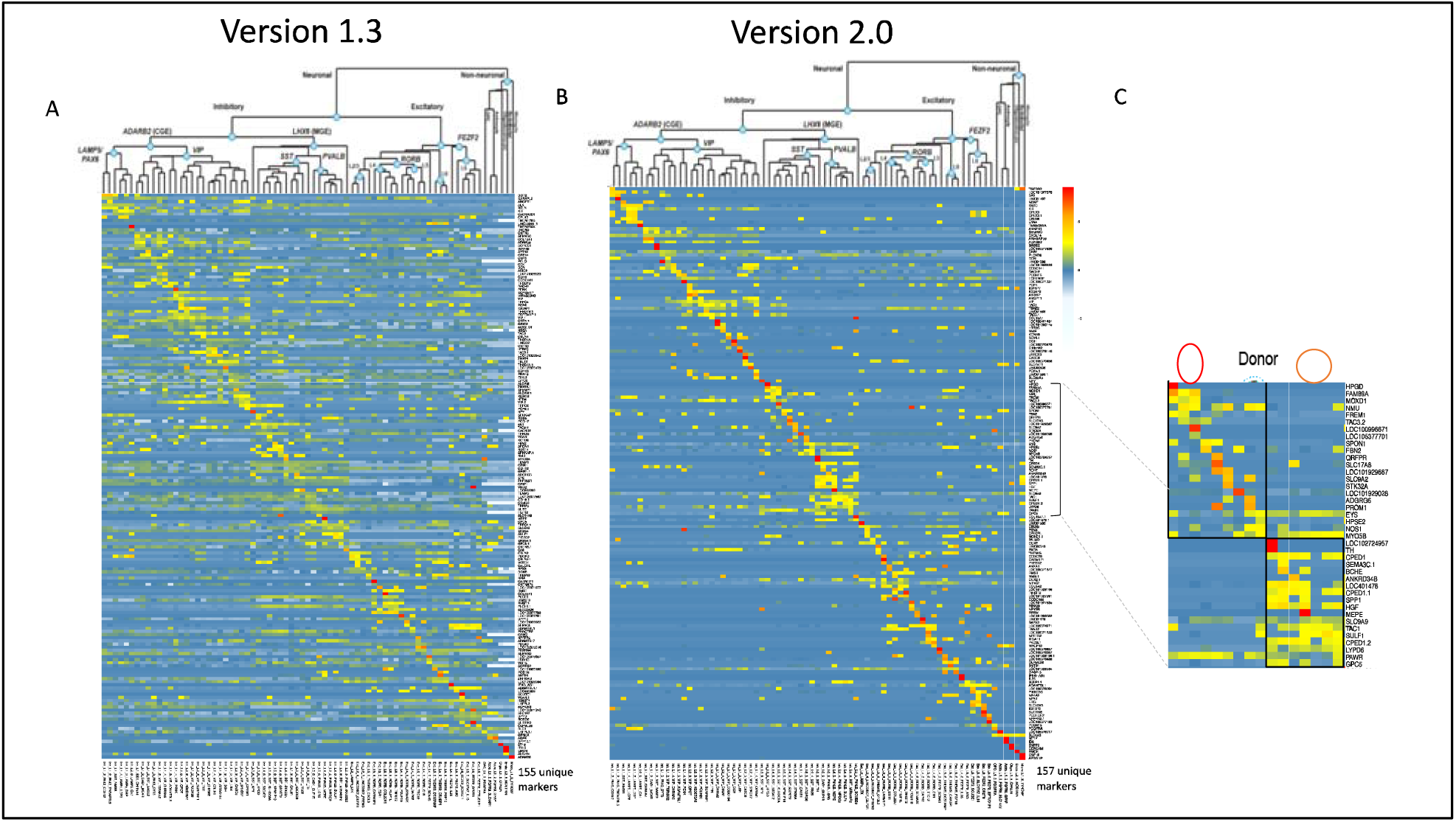
A & B) Heatmaps of NS-Forest v1.3 and v2.0 markers from Human Middle Temporal Gyrus. Clustering solution is described in (citation). Expression values are log2 cluster median normalized by row. Panel C) we have the SST and PVALB clades which demonstrate how relatedness can impact the ability to discover binary markers.

Given the intention of the Binary Expression Score ranking step to preferentially find marker genes with binary expression, there are tradeoffs in both the number of genes required and the classification power when compared to markers ranked by importance from the random forest model. In general, NS-Forest version 2.0 requires more unique genes for a given dataset. In the case of the full MTG dataset the increase is marginal requiring only two additional unique genes (155 vs 157 genes); however, a larger difference in the number of marker genes required has been observed for other datasets (data not shown). Furthermore, the genes that have a high Binary Expression Score are usually not the same genes that were ranked highest by Information Gain within the random forest model. This suggests that in terms of pure classification the markers determined by v2.0 might be expected to underperform. To directly compare the F-scores between these two versions of NS-Forest, an additional analysis was run setting the beta weight of the F-score to 1 in version 2.0 thereby making it directly comparable to version 1.3. As expected, the mean F-score for version 1.3 (0.62) was slightly higher than the mean F-score for version 2.0 (0.58); however, the average Binary Expression Score for the version 1.3 markers was significantly lower at 0.72 versus 0.94 for version 2.0. (For cluster-by-cluster correlations of F-scores and Binary Expression Scores see **sFigure 2**).

Within the major branches of the taxonomy, major subclasses are labeled by important neurological markers such as VIP, SST, and PVALB within the inhibitory subclass, and RORB and FEZF2 within the excitatory subclass. Binary marker genes for the cell types within these subclass branches of the taxonomy can be more difficult to determine, especially when there are many closely related types. For example, both the SST and PVALB contain a number of closely related cell types. When looking at the expression of the marker genes, we can see that between these two major subclasses there is little expression overlap; however, within each subclass there are number of closely-related cell types that show overlapping expression, for example the cell types circled in red within the SST subclass or the types circled in blue within the PVALB subclass (**Figure 3C)**. These cell types tend to have lower F-beta scores and lower marker Binary Expression Scores.

Looking in more detail at the properties of the marker genes selected for individual cell types, we can clearly see the differences between NS-Forest Version 1.3 and 2.0. The expression patterns for the astrocyte cell type Astro_L1_6_FGFR3_SLC14A1 show these differences in the marker genes selected by the two NS-Forest version in more detail (**Figure 4A/B**). NS-Forest v1.3 selects a single marker gene to best discriminate this cluster, while v2.0 selects two. NS-Forest v1.3 selects only the GPM6A gene which performs well at classifying this cell type along a quantitative boundary at the high log2 expression level of 12.5, but also shows intermediate expression in several off-target clusters centered around 10 (**Figure 4A**). Consequently, this quantitative marker is good for classification only when this small window of expression difference is discernible. In contrast, version 2.0 selects LOC105376917 and SLC1A3, both of which have binary expression patterns across clusters (**Figure 4B**). LOC105376917 is highly expressed in only the target cluster and one additional closely-related off-target cluster. Adding SLC1A3 further improves classification by removing cells from this off-target cluster.

**Figure 4:**
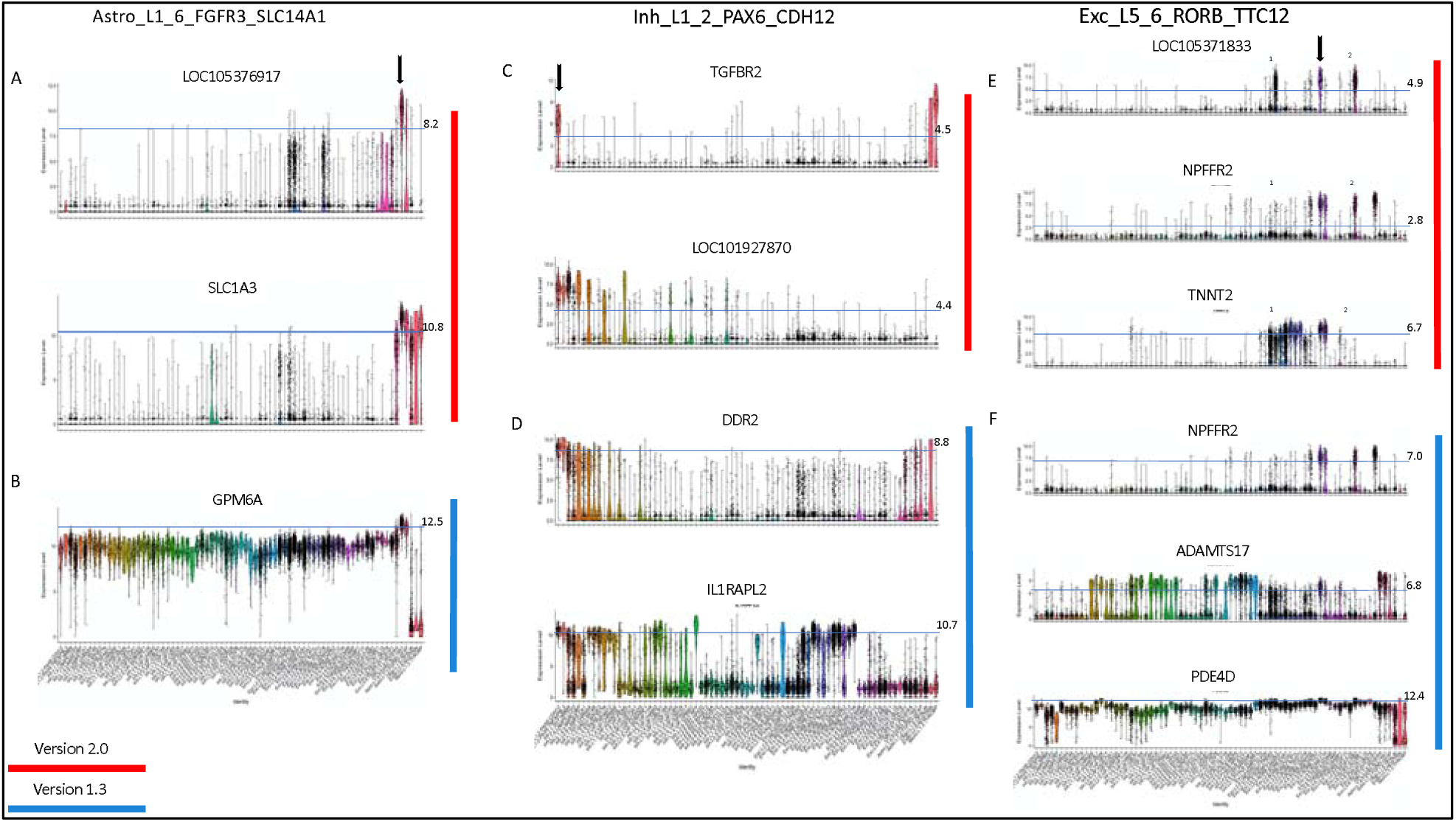
Violin plots for marker gene expression for a selection of cell type clusters representative of the three major classes in the taxonomy: glial cells, inhibitory neurons, and excitatory neurons. Panels A, C, and E (red) give markers determined by NS-Forest v2.0 while panels B, D, and F (blue) give markers from NS-Forest v1.3. Expression is given in log2 scale. Expression thresholds are demarcated by light blue lines and values are given on the right. Thresholds for NS-Forest v2.0 were determined by decision tree split points, while NS-Forest v1.3 were fixed at the cluster median expression for that gene.

In the case of the inhibitory neuron Inh_L1_2_PAX6_CDH12 both v1.3 and v2.0 select two marker genes; however, their characteristics are very different (**Figure 4C/D**). NS-Forest v1.3 again found markers that classified along quantitative boundaries. DDR2 is expressed in all the related clusters in the taxonomy and in some glial clusters at the far end of the taxonomy. The addition of IL1RAPL2 removes the glial clusters and improves the classification; however, IL1RAPL2 is another example of a quantitative marker as it separates the target cluster from the related cluster by narrow differences in expression. NS-Forest v2.0 selected two highly binary markers: TGFBR2, which is very specific to only two clusters, the target cluster and a non-neuronal type at the other end of the taxonomy. The addition of the LOC101927870 gene eliminates cells in the non-neuronal cluster to refine the classification.

Lastly, the excitatory neuron Exc_L5_6_RORB_TTC12 required three markers by both NS-Forest versions to optimize the classification (**Figure 4E/F**). Again, as previously described, NS-Forest v1.3 determined genes that used a quantitative boundary for classification while NS-Forest v2.0 discovered binary markers. A more detailed look at these binary markers provides a clear demonstration of the combinatorics employed by NS-Forest v2.0. Within the target cluster, demarcated by the arrow, all three markers have high expression; however, the off-target excitatory clusters marked as 1 and 2 also express some but not all these markers. By leveraging the combinatorics of the three marker combination, highly discriminative solution is obtained. Gene LOC105371833 is the most binary marker; however, it has high expression in a number of off-target cells in clusters 1 and 2. The addition of the NPFFR2 gene removes most of the false positives in cluster 1, while adding the TNNT2 gene removes the false positives from cluster 2. Together this combination of three marker genes discriminates Exc_L5_6_RORB_TTC12 from other excitatory cell types.

These results show that while adding the Binary Expression Score criteria does slightly decrease the overall classification power of the markers selected, it dramatically increases the binary expression pattern making the markers more useful for downstream applications.

### Characterization of NS-Forest v2.0 Markers

Overall, the results from NS-Forest v2.0 reflect the high quality of the data and clustering analysis as NS-Forest is a supervised machine learning method and is reliant on the quality of the clustering results. The median number of markers required for optimal classification was 2, with only two clusters needing 4 markers, producing a mean F-beta score of 0.69. Overall, the 75 clusters required 157 unique genes to achieve optimal classification. Occasionally, marker genes are shared between clusters, with eleven genes that were not unique [MOXD1, MME, LOC101928196, SULF1, NPFFR2, LINC01583, TAC1, COL15A1, LOC401478, CPED1, TAC3].

Out of the 157 NS-Forest v2.0 marker genes, 37 (24%) were long non-coding RNAs (lncRNAs) or uncharacterized loci (LOCs). Non-coding RNAs have been previously found to be prevalent when analyzing RNA-seq data from single neuronal cells or nuclei and surprisingly these non-coding RNAs had higher specificity as markers when compared to coding genes (27). In particular, lncRNAs are known to show cell line specific expression (28). In contrast, little is known about the LOC genes. These genes are particularly intriguing as they are highly specific to individual cell types and are likely important for their function. As such, they represent areas of unknown biology discovered by scRNA-seq and NS-Forest machine learning that warrant further investigation.

For the characterized marker genes, the most enriched annotations both by adjusted p-value and number of genes involved are for signaling (signal peptide, Signal, GO:0007218∼neuropeptide) and extracellular matrix (Glycoprotein, Extracellular matrix, GO:0005615∼extracellular space, GO:0005578∼proteinaceous extracellular matrix, GO:0030198∼extracellular matrix organization, GO:0005576∼extracellular region, GO:0031012∼extracellular matrix), including neuropeptide, GO:0007218∼neuropeptide signaling pathway, and calcium (**sTable 4**). There are fewer genes annotated with these specific functions as neurology is a rapidly expanding field; however, many other genes assessed here are known signaling peptides in other contexts and would benefit from further characterization in neurological context. Taken together, these results suggests that specific signaling pathways and extracellular signaling molecules are key to neuronal cell type identity.

### Comparison with other Marker Gene Selection Approaches

To understand how the NS-Forest marker genes compare to previously published markers for the human middle temporal gyrus (MTG), we compared the NS-Forest markers to those determined in Hodge *et al* (25) using a different binary expression approach for use in cell cluster naming. In addition to a broad marker determined by the taxonomy and prior knowledge (such as GAD1 or SST), a single marker per cell type cluster was assigned. In total, sixteen of the seventy-five Hodge markers overlapped with the NS-Forest markers [BAGE2, GGH, CASC6, NPY, HPGD, STK32A, ADGRG6, TH, MEPE, PENK, CARM1P1, TWIST2, IL26, SULF1, ADAMTSL1, PDGFRA]. These sixteen were spread across the taxonomy, representing cell type clusters from all three major cell type lineages. Unscaled heatmaps of mean gene expression per cluster for both the Hodge and NS-Forest marker sets (**Figure 5A)** demonstrate that both are characterized by binary expression patterns, having a higher expression along the diagonal versus off-diagonal; however, the Hodge markers have an overall lower mean expression level of 4.8 log2 CPM in comparison with the mean expression for the NS-Forest markers of 7.0 log2 CPM.

**Figure 5:**
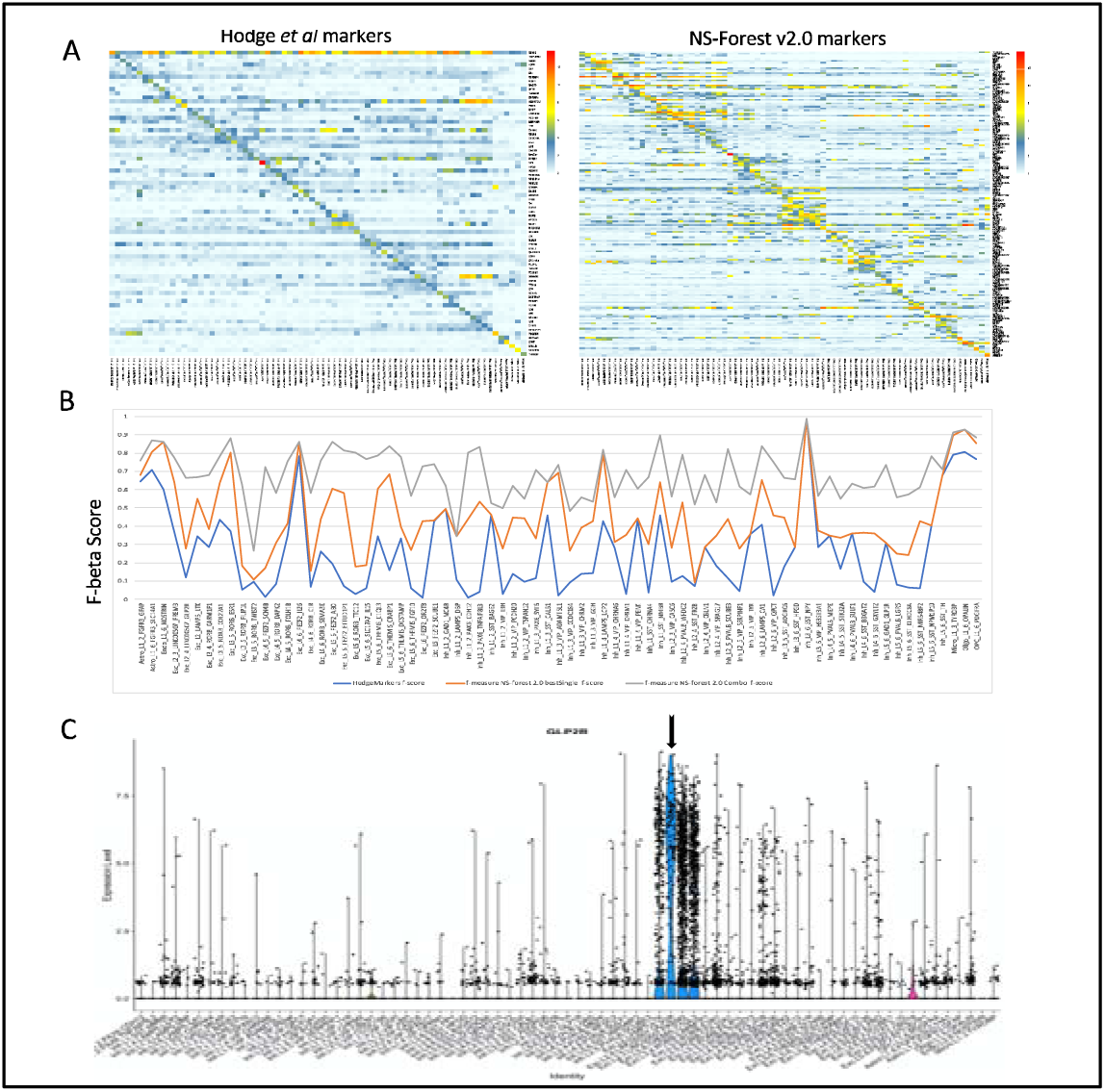
Comparison of Hodge *et al* markers to NS-Forest v2.0 for the full MTG data. Panel A gives an unscaled heatmap where the rows are the mean expression per gene and the columns are clusters. Panel B) Gives the F-beta scores for the single Hodge marker, the best NS-Forest single marker, and the combination of markers found by NS-Forest. C) an example violin plot of a binary expression pattern selected for by the method used by Hodge *et al* for cluster Exc_L2_4_LINC00507_GLP2R.

One major difference between these two approaches is that the Hodge marker set contains a single marker per cluster to label a distinct cluster phenotype while NS-Forest selects combinations of markers that optimize classification power. By running the Hodge markers through NS-forest v2.0, we estimated F-beta scores for the single Hodge markers in order to compare their classification power to the best single NS-Forest markers, and the NS-Forest combination of markers (**Figure 5B)**. Overall, the trend lines show that the F-beta scores for single markers, both blue and orange lines, follow a similar trajectory with some clusters being more difficult to classify then others, i.e., having lower F-beta scores. However, the NS-Forest combination of markers, shown in grey, demonstrate that combinations of markers yield a uniformly higher power of discrimination over a single marker, regardless of how the single best marker is chosen.

When evaluating the F-beta scores for the Hodge markers, it became clear that many had elevated false positive rates. To directly compare the two sets of markers, we computed the false discovery rate (FDR= FP/FP+TP) for each cell type and averaged across the entire set. The Hodge markers had an average FDR of 0.7 versus 0.14 for the NS-Forest markers. GLP2R, which is a marker for Exc_L2_4_LINC00507_GLP2R, offers a good visual example (**Figure 5C)**. This gene expressed in the target cluster but also the nearest cell types within the LINC00507 group. NS-Forest also has difficulty finding markers for this cell cluster phenotype, requiring 3 markers in total; however, in combination these markers help reduced the FDR rate from 0.89 to 0.11.

### Validation of Human MTG NS-Forest v2.0 Markers

At current, the ground truth for the neuron types and their marker genes in human MTG taxonomy is not available as it is an active area of investigation. Consequently, a true biological validation of the marker genes is not possible. As an alternative, we asked the question, do the minimum set of marker genes selected by NS-Forest capture the underlying structure of cell type identity reflected in the entire expressed transcriptome? To do this, we generated tSNE plots using the complete 5574 variable genes used for the original MTG clustering, the minimum set of 157 NS-Forest v2.0 marker genes, and 157 genes randomly selected from the complete variable genes list. These embeddings were then painted using the cell type assignments from the MTG taxonomy. From the tSNE plots it is clear that the NS-Forest markers closely recapitulate the clustering structure of the complete variable genes set, much better than the randomly selected genes (**Figure 6A)**. For example, in the bottom of the complete variable gene tSNE there is a light salmon and dark salmon colored group of clusters, and these two clusters are preserved in the right hand side of the NS-Forest marker tSNE, whereas in the tSNE from the randomly selected variable genes these two clusters spread out and a third brown cluster is now merged with light salmon cluster. Examples like this can be seen throughout the three embeddings. A more quantitative analysis of these tSNE embeddings using the Nearest-Neighbor Preservation metric shows that both the precision and recall are higher using the 157 NS-Forest markers compared with 50 sampling of 157 genes randomly selected from the variable gene set (**sFigure 3**).

**Figure 6:**
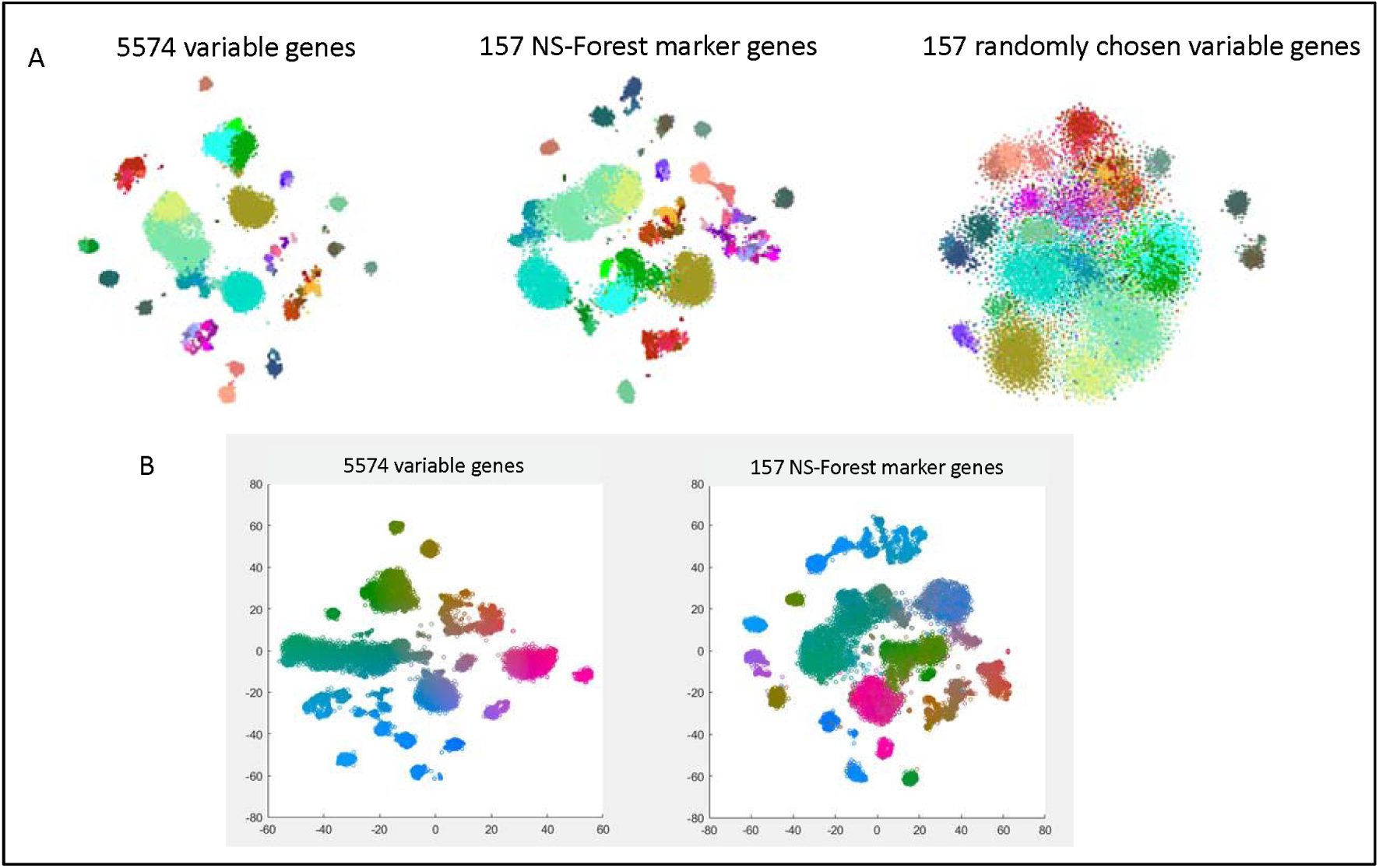
Validation of the 157 NS-Forest v2.0 Middle Temporal Gyrus (MTG) marker genes. In panel A) we have tSNE plots for the full DE list of 5574 genes, the 157 NS-Forest markers, and 157 genes randomly selected from the variable gene list. In panel B) left is a tSNE generated from the full variable gene list of 5574 genes colored by coordinate position while right is a tSNE generated using the 157 NS-Forest markers then painted by nuclei according to the color scheme established in the full tSNE on the left.

In addition, the local embedding structures within a given tSNE cluster also appear to be well preserved (**Figure 6B)**. The complete variable gene tSNE was painted using a color gradient based on the coordinate positioning. This yields a visual way of comparing where individual nuclei are located within the full tSNE embedding versus other tSNE embeddings. The NS-Forest marker tSNE was then painted using the colors derived from the full tSNE gradient. The fact that the same color gradients are observed in the NS-Forest embedding demonstrates that the positional gradients, and thus the nuclei-to-nuclei relationships, in the NS-Forest embedding closely reflect the positional gradients in the complete full tSNE embedding. For example, in the full tSNE there is a long cluster of nuclei beginning on the left in green that extends toward the middle moving into bluish green and ending with a purplish blue. This same cluster, with the same color gradient is preserved within the center left cluster of the NS-Forest tSNE.

## Discussion

Here we describe NS-Forest version 2.0. Development was driven by user community requirements for data derived cell type phenotype definitions that are testable in future investigations. To this end, a number of changes were made after the random forest feature selection. In earlier version of NS-Forest, negative markers were occasionally found. These are marker genes that are expressed in most off-target clusters but not the target cluster. Given that testing for a something that is not expressed is methodologically difficult, it was decided to avoid this category of markers. By implementing a median expression level cutoff greater than zero for the target cluster, we removed all possible negative marker genes. In addition, this cutoff also defines one basic characteristic of a NS-Forest Marker: they are required to be expressed in greater than half of the cells within the cell type cluster.

NS-Forest v1.3 contained simple random forest feature selection approach that discovered quantitative markers that were good for classification but generally problematic for further biological investigation. This limitation of random forest feature selection may be shared with other machine learning methods. Consequently, a ranking step was incorporated to select for markers with binary expression patterns. Simulation testing performed on this Binary Expression Score ranking step demonstrated that it selected for marker genes with binary expression patterns and accurately ranked them according to level of binary expression. As a result, NS-Forest v2.0 demonstrated clear improvement in the enrichment for binary expression patterns but at a small cost to the overall classification power and number of marker genes necessary. Consequently, If the user requires classification with less requirement for downstream investigation, then we would recommend using NS-Forest v1.3; however, in all other cases NS-Forest v2.0 is recommended. Both versions are available as official releases at the github repository.

Beyond their use for defining and investigating cell type phenotypes, necessary and sufficient marker genes also offer a dimensionality reduction with limited loss of fidelity to the originally clustering solution. This dimensionality reduction offers a feasible way of representing the clustering solution with a minimal amount of information which is ideal for data dissemination. These marker genes can then be used to generate a reference knowledgebase for cell types, in effect generating an expression barcode of marker genes for a given cell phenotype.

As mentioned above, NS-Forest markers are optimized for downstream experimental investigation. There are a number of assays for which known markers could facilitate biological investigation such qPCR and the burgeoning field of spatial transcriptomics based on multiplex FISH. To date a number of projects have used NS-Forest markers for these purposes. For example, qPCR probes based on NS-Forest markers were made to detect genes in scRNA-seq libraries from myeloid dendritic cells (mDCs) FACS sorted from peripheral blood in patients treated with the Hepatitis B vaccine (30, publication in preparation). In a similar fashion, gene probes were designed based on NS-Forest markers for cell type detection using a number of spatial transcriptomic technologies. These technologies aim to resolve the location within the tissue of cell types derived from scRNA-seq generated taxonomies (31).

Another possible application of NS-Forest is to utilize selected gene sets of particular interest as input to produce marker gene sets designed to capture specific cell type properties. For example, the input of gene sets composed of transcription factors could reveal master regulators of developmental programs (32). Input gene sets composed of neuropeptides and neurotransmitters could be used to shed new light on the specific signaling properties of different neuronal cell subsets (33). Input gene sets composed of cell surface markers could be used to identify markers for use in FACS sorting.

As the number of experiments performed and datasets made publicly available dramatically increase, the greater biological community is left with the monumental task of integrating these data into a consensus of canonical cell types. With cell phenotypes defined by NS-Forest marker genes, we can move ahead with the creation of a dissemination framework that defines ontological classes based upon these molecular markers as the necessary and sufficient criteria in an axiomatic semantic representation compliant with FAIR principles. Ontological representation has numerous advantages over simple vocabularies, including the structuring of knowledge in a computationally readable format so that findings from many experiments can be easily accessible and “reasoning” can be performed to ensure the consistency of the representation as the knowledge rapidly grows. These provisional instances of “cell type clusters” defined by NS-Forest markers can form the basis for the instantiation of an ontology class that can be in the future adopted into the official Cell Ontology (CL). Progress is already underway in developing programmatic and scalable methods to handle the amount of single cell data being generated. This ontological representation can address several pressing needs of the wider biological research community. Producing an easy, visually accessible overview of the results of many single cell experiments in a traversable structure while preserving the hierarchical relationships inherent in a taxonomy of cells. In addition, this ontology will provide a platform for integration with other data modalities such as cell morphology, electrophysiology, cell-cell interactions. A provisional cell ontology (pCL) generated in this manner for Middle Temporal Gyrus and primary motor cortex is available for exploration at https://bioportal.bioontology.org/ontologies/PCL.

## Methods

### NS-Forest version 2.0

Initial Feature Selection: The NS-Forest version 2.0 workflow (**Figure 1a-b**) begins with a cell-by-gene expression matrix, with an additional column containing cluster membership labels, produced by any expression data clustering method applied to single cell/nucleus RNA sequencing (scRNA-seq) datasets. This cluster-labelled expression matrix is then used to generate Random Forest classification models distinguishing each target cluster from all other clusters (binary classification) using RandomForestClassifier scikit. RandomForestClassifier hyperparameters were left at default except that the number of trees was set at 10,000 to give sufficient coverage of the sample and gene expression feature space; necessary coverage for a given feature space is estimated as the square root of the number of samples (∼10,000 cells) times the square root of the number of features (∼10,000 genes). From the resulting Random Forest model, the average Gini Impurity value is used to initially rank genes based on their feature importance.

#### Feature Re-ranking Based on Positive Binary Expression

Re-ranking the features after initial Random Forest selection begins with positive expression filtering (**Figure 1c**). By default, genes with a median cluster expression of 0 are removed in order to exclude genes that are not expressed in the relevant cluster, which we refer to as negative markers, or show high zero inflation. This parameter is tunable and can be adjusted according to the desired positive expression level.

Next, genes are re-ranked to enrich for genes with binary expression patterns (**Figure 1d)**. A “Binary Expression Score” was developed to select for genes that show all-or-none expression patterns, with expression in the target cluster and as few other cell type clusters as possible. The Binary Expression Score is calculated for each gene in the initial Random Forest feature list according to the equation:

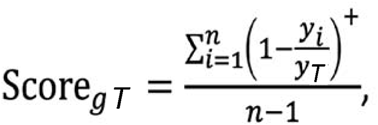

where y_i_ is the median gene expression level for each cluster i, y_T_ is the median expression in the target cluster, and n is the number of clusters. This results in a Binary Expression Score in the range of 0 – 1, with a Binary Expression Score of 1 being the ideal case where the gene is only expressed in the target cluster (**Figure 1e**).

#### Minimum Feature Combination Determination

After the top genes are re-ranked based on positive binary expression, they are then tested for their classification power individually and in combination. First, the top M genes (6 genes by default) are used to generate individual decision trees to determine the optimal expression level cut-off value for each gene (**Figure 1F**). The maximum leaf nodes parameter is set at two, thereby ensuring a single split point per tree. From these trees, the optimal gene expression threshold at the split point is extracted.

To evaluate the discriminative power of a given combination of candidate marker genes, we use the F-beta score as an objective function. The F-score is the harmonic mean of precision and recall providing equal weight for these two classification measures. The F-beta score includes a beta term that allows for the weighting of the function towards either precision (beta < 1) or recall (beta > 1) (**Figure 1G**). The beta for the analysis described here was estimated empirically at 0.5 (**Supplemental Figure 1**).

Finally, all combinations of the top ranked genes (6 genes by default) are then evaluated at the expression levels determined earlier by decision tree analysis. The F-beta scores for all combinations are written to a complete results file and the gene feature combination producing the best F-beta result selected per cluster.

### Simulation Testing of the Binary Expression Score

Simulation studies were conducted to investigate the properties of the Binary Expression Score weighting using a three-component mixture model to reflect the zero-inflation technical artifact and the background and positive expression signals in real data distributions. Denoting X as the gene expression value, our simulated data follow a mixture distribution:

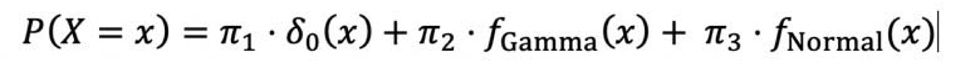

Where δ_0_(x) is the probability density function of the degenerate distribution at 0 for the zero-inflation technical artifact, f_Gamma_(x) is the probability density function of a Gamma distribution (with hyperparameters α and β) for low level background expression from off-target cells or on-target cells with low expression, and f_Normal_(x) is the probability density function of a Normal distribution (with hyperparameters μ and σ) for positive expression signals; parameters π_1_, π_2_ and π_3_ are the corresponding mixture weights for each component such that π_1_, π_2_, π_3_ > 0 and π_1_+ π_2_+π_3_ = 1. In our simulations, we generated 20 clusters with 300 cells in each cluster. We designed cases where the simulated gene is expressed at high levels in 1, 2, or 5 clusters. Both binary and quantitative markers were simulated for on-target and off-target clusters by setting different parameters and hyperparameters in the mixture model.

### snRNA-seq Data

The scRNA-seq data evaluated here were obtained from the Allen Institute for Brain Science (https://portal.brain-map.org/atlases-and-data/rnaseq). Experimental design, including tissue sampling and data processing, can be found in Hodge *et al*. (23). In brief, layers 1-6 of the human Middle Temporal Gyrus (MTG) were vibratome sectioned, nuclei were extracted and labelled for NeuN expression. Nuclei were then FACS sorted and libraries were generated using the Smart-Seq4 and Nextera XT chemistries. Data processing and clustering were then performed as detailed in (22).

NS-Forest v2.0 was run using the cluster assignments given in Hodge *et al*. (23). Cells not assigned to a cluster were removed from the analysis. CPM expression values were log2 (x+1) transformed and genes with a sum of zero median expression across all clusters were removed. After filtering, 15,928 cells and 13,946 genes remained. Given the size of the input matrix, we increased the number of trees in the random forest model from the default of ten thousand to fifty thousand.

### Marker Validation

In order to demonstrate the preservation of the cell type clustering characteristics using NS-Forest marker genes, tSNE embeddings were generated using Cytosplore. The original clustering solution is represented by an embedding generated from the 5574 variable genes used for the iterative clustering originally performed (22, 23). Additional embeddings were made using the combined set of 157 marker genes for all cell type clusters determined by NS-Forest version 2.0, and 50 sets of 157 genes chosen at random from the original 5574 genes.

## Data Access

All data used herein is publicly available.

## Acknowledgements

This work was supported by the Allen Institute for Brain Science, the JCVI Innovation Fund, the U.S. National Institutes of Health (R21-AI122100 and U19-AI118626), the California Institute for Regenerative Medicine (GC1R-06673-B), the Wellcome Trust 208379/Z/17/Z, and from the Chan Zuckerberg Initiative DAF, an advised fund of the Silicon Valley Community Foundation (2018-182730).

## Supplemental Figures (legends only)

**sFigure 1:**
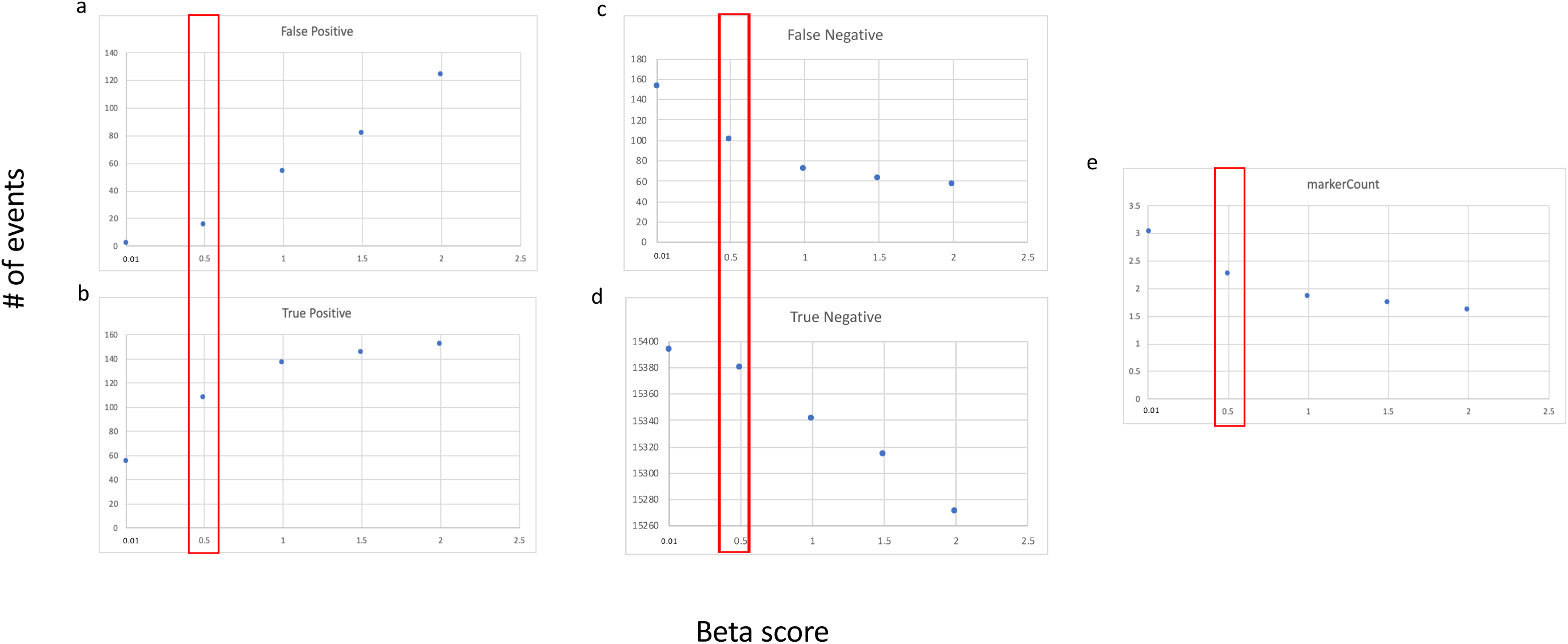
Empirical determination of the beta weighting parameter for the F-beta score. All 75 clusters from full Middle Temporal Gyrus data were used to estimate the average true positive (TP), false positive (FP), false negative (FN), and true negative (TN), and number of markers for all clusters at beta values of 0.01, 0.5, 1, 1.5 and 2.

**sFigure 2:**
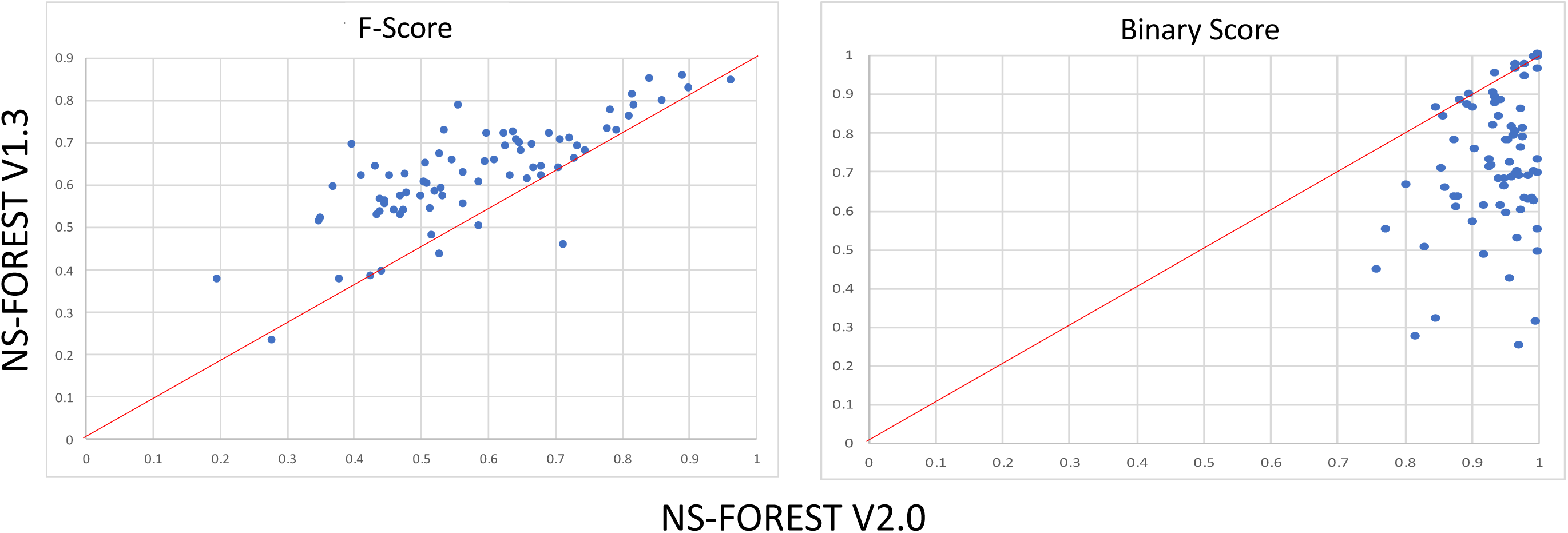
Correlation between F-scores and average Binary Expression Scores per cluster for v1.3 and v2.0. The F-score per cluster was computed using a beta=1 for NS-Forest v2.0 to make both versions comparable.

**sFigure 3:**
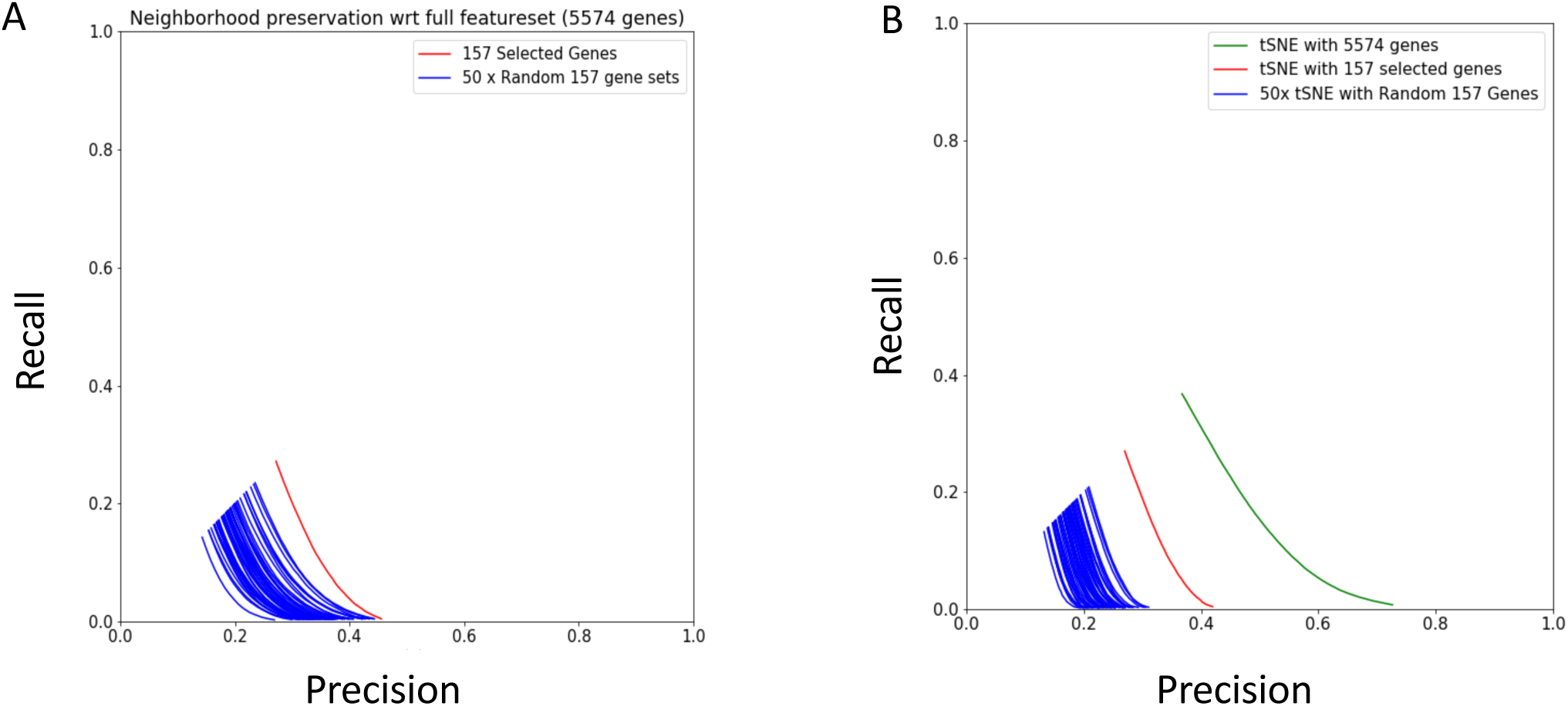
Quantitative assessment of Nearest-Neighbor Preservation metric (NNP, by Venna et al. and IM). In brief, this is computed as follows: for each data point, the K-Nearest-Neighborhood (KNN) in the high-dimensional space is compared with the KNN in the reduced-dimensional space. Average precision/recall curves are generated by taking into account high-dimensional neighborhoods of increasing size up to Kmax = 50. The True-Positive number is the intersection between high-dimensional and the low dimensional neighborhood based on 157 selected genes. The precision is computed as TP/K and the recall as TP/Kmax. In A) red curve: NNP curves for the random-forest selected 157 genes, while blue curves: NNP curves for 50 random gene sets of the same size (selected from the full 5574 high-variance gene set). In B) green curve is the tSNE generated from the complete 5574 variable genes, the red curve: NNP curves for the random-forest selected 157 genes, while blue curves: NNP curves for 50 random gene sets of the same size (selected from the full 5574 high-variance gene set)

sTable 1: Complete NS-Forest results for v1.3

sTable 2: Complete NS-Forest results for v2.0

sTable 3: Supplemental ranked binary results from NS-Forest v2.0

sTable 4: Enrichment of annotations

